# Implications for human odor sensing revealed from the statistics of odorant-receptor interactions

**DOI:** 10.1101/283010

**Authors:** Ji Hyun Bak, Seogjoo J. Jang, Changbong Hyeon

**Affiliations:** School of Computational Sciences, Korea Institute for Advanced Study, Seoul, Korea; Department of Chemistry and Biochemistry, Queens College, City University of New York, New York 11367, USA; PhD programs in Chemistry and Physics, and Initiative for Theoretical Sciences, Graduate Center, City University of New York, New York 10016, USA

## Abstract

Binding of odorants to olfactory receptors (ORs) elicits downstream chemical and neural signals, which are further processed to odor perception in the brain. Recently, Mainland *et al*. [*Sci. data*, (2015) 2:sdata20152] have measured ≳ 500 pairs of odorant-OR interaction by a high-throughput screening assay method, opening a new avenue to understanding the principles of human odor coding. Here, using a recently developed minimal model for OR activation kinetics [*J. Phys. Chem. B* (2017) 121, 1304-1311], we characterize the statistics of OR activation by odorants in terms of three empirical parameters: the half-maximum effective concentration EC50, the efficacy, and the basal activity. While the data size of odorants is still limited, the statistics offer meaningful information on the breadth and optimality of the tuning of human ORs to odorants, and allow us to relate the three parameters with the microscopic rate constants and binding affinities that define the OR activation kinetics. Despite the stochastic nature of the response expected at individual OR-odorant level, we assess that the confluence of signals in a neuron released from the multitude of ORs is effectively free of noise and deterministic with respect to changes in odorant concentration. Thus, setting a threshold to the fraction of activated OR copy number for neural spiking binarizes the electrophysiological signal of olfactory sensory neuron, thereby making an information theoretic approach a viable tool in studying the principles of odor perception.

## Introduction

Olfaction, ubiquitous among all animals, is a key sensory process that is used to detect a vast number of chemicals in the external world [1, 2]. From the perspective of molecular recognition, the physicochemical principle germane to the early layer of olfactory process is not significantly different from that of unicellular organisms’ chemotactic response [3, 4, 5]. Similarly, olfactory signals are initiated upon the recognition of odorants by the receptors that are expressed in the nasal epithelium [6].

Physiological and biochemical studies of olfaction to date offer strong evidence that the majority of mammalian OR signalings are associated with G-protein dependent pathway [7]. Thus, except for a few cases [8, 9, 10], their activation mechanism is mostly related to that of G-protein coupled receptors (GPCRs) [11, 12, 13, 14, 15]. Upon binding odorants, a set of ORs adopt their structures into active forms and catalyze G-proteins to initiate downstream signal cascades. Although the original input signal is modified passing through multiple layers of neural circuits [16, 17], elucidating the information encoded to the pool of ORs is a key component for understanding the principles of odor sensing. In particular, the receptor code, or the responses of a repertoire of ORs encoding the chemical features of odorant(s), constitutes the first layer of dataset to be mapped by the olfactory process.

Unlike the conventional GPCRs that are deemed ‘fine-tuned’ to endogenous agonists such as hormones and neurotransmitters [18, 19, 20, 14], ORs tend to be ‘broadly tuned’ to multiple odorant types. Conversely, a single odorant can also be recognized by multiple ORs. In fact, there is increasing evidence indicating that even the conventional GPCRs can adopt multiple active states and yield different signaling pathways [21, 22], replacing the old view that GPCR activation is described by the two-state conformational selection model [23]. The many-to-many interactions of ORs with odorants enable a limited number of ORs to efficiently encode a huge chemical space represented by the odorants and their mixtures [24, 25]. The human olfactory system is known to employ *N_r_* ≈ 330 OR subtypes [26], whereas the odors are made of an estimated pool of *M* ≳ 10^4^ monomolecular odorants [27, 28] and their *m*-component mixtures. To be able to discriminate potentially a vast number of natural odors 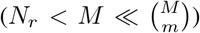, the olfactory system adopts a strategy based on *combinatorial coding* [29]. In a simplifying limit of ON/OFF switch-like response, a set of *N_r_* receptor types can, in principle, generate 2^*N_r_*^ ≈ 2^330^ ≃ 10^99^ distinct receptor codes, which allow human neural systems to discriminate *M^m^* distinct natural odors for fairly large value of *m*.

Traditionally, the odor perception has been interrogated by measuring the brain activity [30] or through psychophysical tests [31]; however the associative and nonlinear nature of neuronal codes [32] and genetic variation among human subjects [33] make it difficult to decipher a direct connection between odorants and the final odor perception. Recent developments of experimental techniques using high-throughput screening [34] have compiled dose-response data from a set of 535 interacting pairs of odorants and ORs. The combinatorial nature of odor coding was substantiated by the concrete, objective dataset demonstrating the responses of 304 human ORs against 89 odorants [34].

The aim of this work is to gain more quantitative insights into the physicochemical processes that take place in the early stage of odor sensing. More specifically, we aim to shed light on the statistics of odorant recognition and infer its relation with a cellular response of the corresponding olfactory receptor neuron (ORN). To this end, we adapt a minimal kinetic model for the activation of mammalian ORs [35], and quantitatively analyze a set of doseresponse data for human ORs [34]. Analyzing the experimental data, we characterize the statistics of odorant-receptor interactions in terms of three empirical parameters - the half-maximum effective concentration (EC_50_), efficacy, and basal activity - and obtain the distribution of these parameter values. In particular, we relate the three parameters with the rate constants and the binding affinities, and specify the condition to be met by the kinetic parameters of OR activity. Because the number of ORs expressed in each ORN is large (*L* ≈ 2.5 × 10^4^ [36]) and the signals from on average 10^4^(≈ 6 × 10^6^/350) ORNs [37] of the same type are converged to a glomerulus [38], the response of ORN can be best understood as an outcome that integrates stochastic OR signals over the population. Resorting to the law of large number and the concept of spike firing threshold, we propose that the early stage of olfactory neural signals can be binarized, making an information theoretic approach as a viable tool for studying the odor sensing.

## Model

### Minimal kinetic model for OR activation

In order to model the OR activation we consider the following *minimal kinetic scheme* [35]:

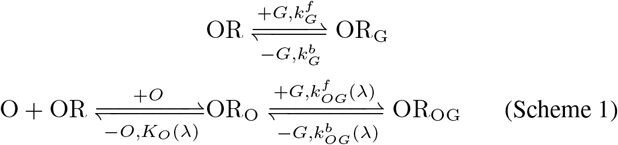

where the first and second lines represent binding/unbinding of the G-protein with the OR in the absence and presence of an odorant (*O*) in its binding site, respectively. 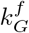 is the rate of G-protein (GDP-bound heterotrimeric complex) binding to OR without odorant, and 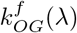 is the rate of G-protein binding when an odorant of the type *λ* is bound to the OR. *C_G_* is the intracellular concentration of G-protein. *K_O_* is the dissociation constant of the odorant from OR that can be related to the odorant-OR binding affinity (*A*) as *A* ≡ Δ*G_diss_* = −*k_B_T* log *K_O_* where Δ*G_diss_* is the dissociation free energy associated with OR_O_ ⇌ O + OR; 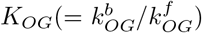 and 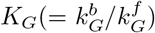 are the dissociation constants of G-protein from OR, with and without odorant in the OR, respectively (see Fig 1a). In Scheme 1, the intracellular down-stream signal cascade is initiated by the kinetic steps marked with a “–*G*”, indicating a G*α*_GTP_ subunit released from a heterotrimeric G-protein complex. Even without an odorant in the binding pocket of OR, OR can relay weak downstream signal cascades which are defined as the basal activity. For an OR that binds a cognate odorant (OR_O_), the binding of G-protein is expected to be further facilitated 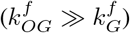, evoking a stronger downstream signal cascade (Fig 1a).

**Figure 1:**
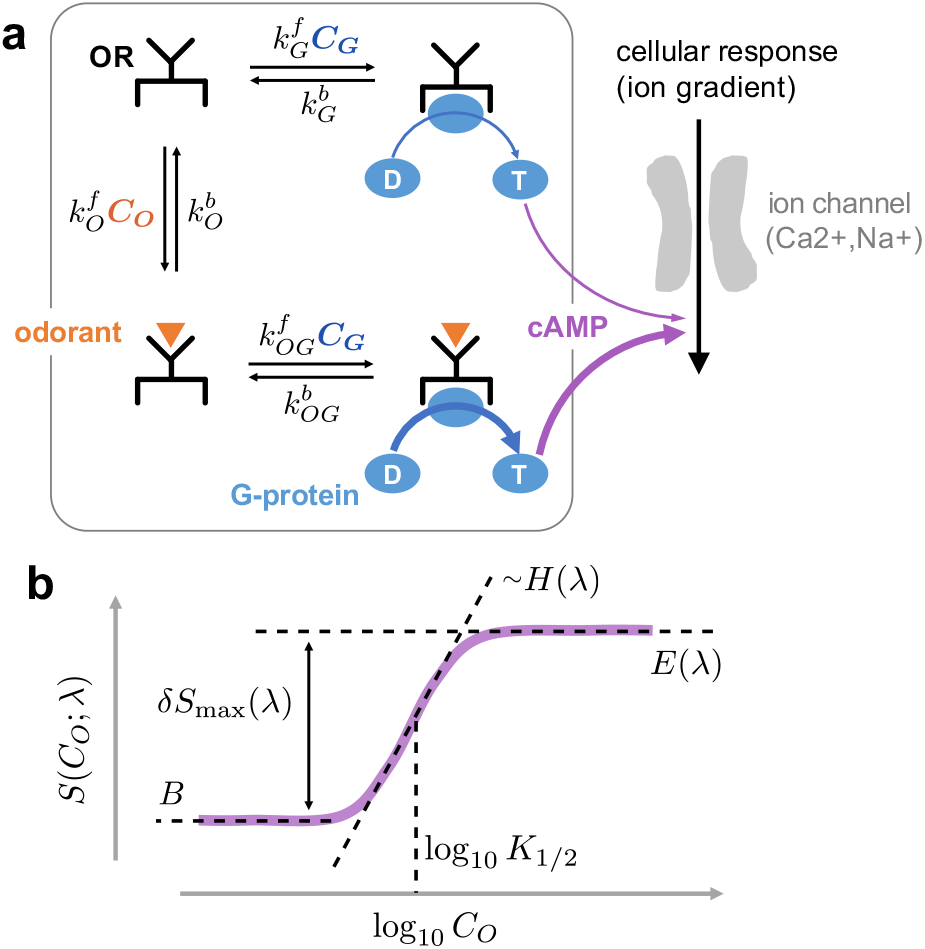
The minimal kinetic model for the odor response. **a**. A schematic of olfactory signaling cascades. G_olf_, olfactory G-protein binding to and the subsequent release of G*α*_GTP_ subunit from the receptor via GDP/GTP exchange 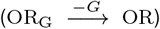 elicit downstream signal cascades: production of second messengers (cAMP) via stimulation of adnenylyl cyclase III, followed by depolarization of membrane potential via cAMP-induced opening of ion-channels [39, 40]. The downstream signal is stronger (the thicker arrow in purple) when odorant is bound to the OR. **b**. The four-parameter model (Eq 3) to describe the dose-response data.

Based on the minimal signaling scheme (Scheme 1), the OR activity *S*(*C_O_*; *λ*), evoked by an odorant of the type *λ*, can be shown to be a hyperbolic function of the odorant concentration *C_O_* [35]:

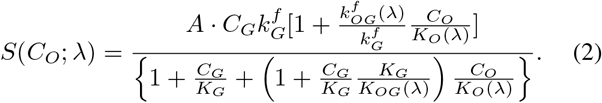

Because the activity of each receptor in Mainland *et al*.’s assays [34] was measured by means of the fluorescence emitted from cAMP-mediated luciferase activity, we assume here that the strength of cellular (ORN) response is proportional to the amount of Gα subunit released from the complex and consequently the cAMP-mediated gene expression. The proportionality constant *A* reflects all of these kinetic details.

### Fitting the dose-response data using the Hill curve

It is useful to cast the above model (Eq 2) into the following Hill equation:

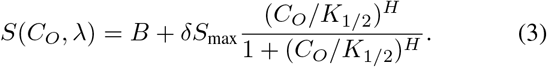

The three key parameters of the dose-response relationship (see Fig 1b), EC_50_[= log_10_ (*K*_1/2_/[*M*])] where *K*_1/2_ is the halfmaximum concentration in molarity unit, the efficacy (*E* = *B* + *δS_max_*), and the basal activity (*B*), are expressed in terms of the rate and dissociation constants defined in the reaction scheme for OR signaling (Scheme 1 and Eq 2):

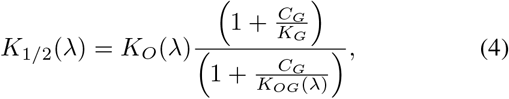

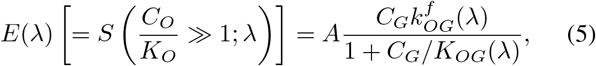

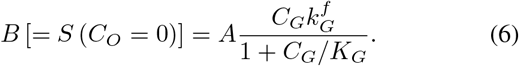

We have specified *λ* in the argument of parameters to make it explicit that the parameters depend on a specific odorant-OR pair. Note that for a given receptor, *K*_1_/_2_ and *E* change with the odorant type *λ*, whereas *B* is a property of the receptor only. Eq 3 is equivalent to Eq 2 when the Hill coefficient is fixed to *H* = 1. Empirically, however, we find that the dose-response data can be best fitted by treating *H* as a free parameter.

## Results

### A universal response curve for human olfactory receptors

We fitted the dose-response data of [34] to the Hill curve partly based on our kinetic model (Eq 3). Among 535 odorant-OR pairs in the dataset, 475 pairs showed activation (*δS*_max_ > 0), and the rest showed deactivation (*δS*_max_ < 0). Furthermore, 317 out of 535 pairs could be fitted to Eq 3 reliably with correlation coefficients greater than 0.9. Fig 2a-d show examples of odorant-OR pairs. By using the parameter values fitted for respective odorant-OR pairs, and using the rescaled variables *ĉ* ≡ (*C_O_*/*K*_1/2_)^*H*^ and *f* ≡ *δS*/*δS*_max_ = (*S − B*)/*δS*_max_ we could collapse the dose-response data on a universal curve, *f* = *ĉ*/(1 + *ĉ*) (Fig 2e). This justifies our use of Eq 3 for the analysis of the odorant-OR dataset.

**Figure 2:**
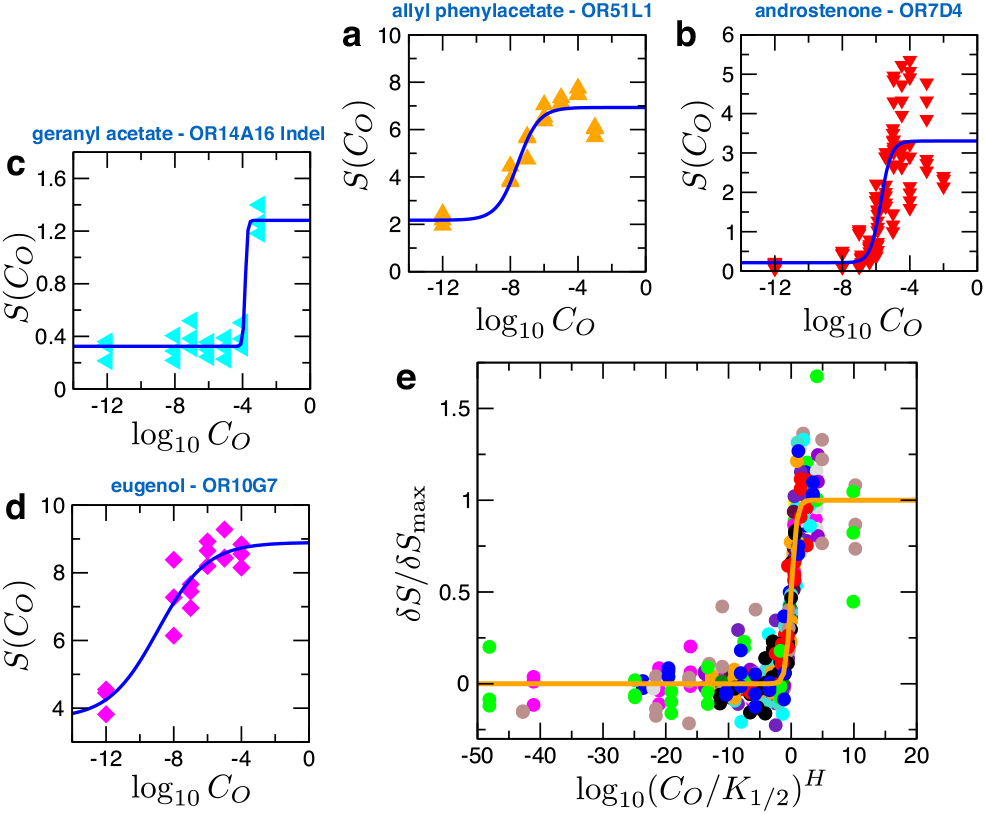
Analysis of dose-response data for odorant-OR pairs. Examples of fit using Eq 3 are shown for some odorant-receptor pairs: **a**. allyl phenylacetate-OR51L1 (*B* = 2.18, *δS_max_* = 4.76, log_10_ *K*_1/2_ = −7.56, *H* = 0.7); **b**. androstenone-OR7D4 (*B* = 0.21, *δS*_max_ = 3.09, log_10_ *K*_1/2_ = −5.72, *H* = 1.4); **c**. geranyl acetate-OR14A16 Indel (*B* = 0.32, *δS*_max_ = 0.96, log_10_ *K*_1/2_ = −3.84, *H* = 6.6); **d**. eugenol-OR10G7 (*B* = 3.69, *δS*_max_ = 5.21, log_10_ *K*_1/2_ = −8.95, *H* = 0.3). **e**. Dose-response data for 22 odorant-OR pairs collapsed onto a universal curve, *f* = 10^*ξ*^/(1 + 10^*ξ*^), where *f* = *δS*/*δS*_max_ and *ξ* = *H* log_10_(*C_O_*/*K*_1/2_) (orange line). For clarity, only the data from 22 activating pairs are presented.

### Distribution of the empirical parameters over the ensemble of receptors

In order to gain a rough statistical estimate of the sensitivity of human ORs we examine the range of odorant concentration to which human ORs are responsive, and the variation in the magnitude of intra-cellular response elicited by OR activation. The odorant-OR interaction dataset [34] allowed us to obtain the distributions of the two key empirical parameters, the effective concentration EC_50_ and the efficacy *E*, over the ensemble of receptors.

First, we constructed the histogram of the effective concentrations of odorants from all the data of interacting odorant-OR pairs (see Fig 3a), and obtained the distribution *ψ*_ens_(EC_50_) by normalizing the histogram, where the subscript “ens” indicates that the distribution was constructed from the entire *ensemble* of odorant-receptor pairs. The concentrations exhibit a fairly broad distribution, ranging from nM (EC_50_ = −9) to M (EC_50_ = 0), with an average 〈EC_50_〉 = −4.1 and a standard deviation of *σ*_EC_50__ = 1.8. This corresponds to a range of 1 *μ*M < *K*_1/2_ < 5 mM. Most of the odorant-OR pairs with strong affinities (EC_50_ < −9) are deactivating pairs, in which the odorants act as the inverse-agonists of the respective ORs (Fig 3a). Compared to the proportion of all the odorant types tested by Mainland *et al*. [34], the odorants contributing to the deactivating pairs with strong affinities mostly belong to certain specific chemical types. They are mainly identified to be acidic (thioglycolic acid, isobutyric acid, 3-methyl-2-hexenoic acid), aromatic (cumarin, quinoline, 2-methoxy-4-methylphenol), acetate (n-amyl acetate, butyl acetate), steroid (androstenone), and ester (pentadecalactone, amyl butyrate) (see Fig S2). Presently, the molecular cause for the inverse agonism by these strongly bound odorants is not clear, but it has been suggested that sidechains of GCPRs interacting with inverse agonist are generally more rigid [41].

**Figure 3:**
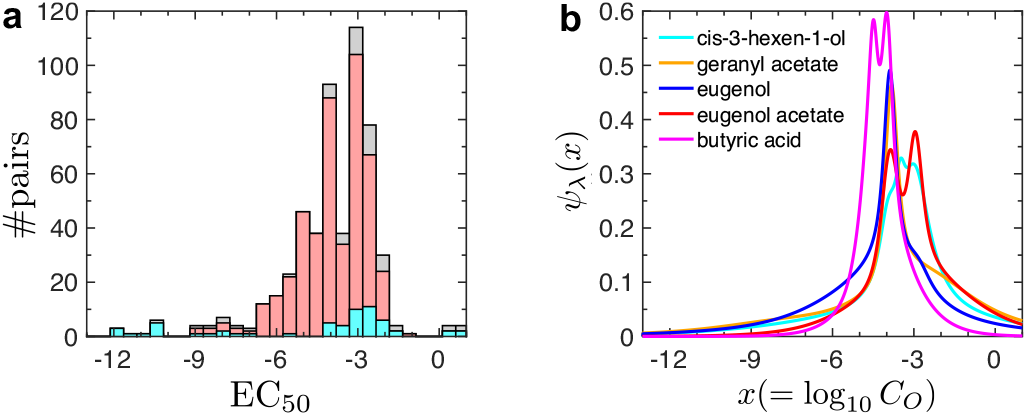
Distribution of effective odorant concentration for the activation of partner receptors. **a**. Histogram of EC_50_ calculated for all dose-response data (gray). Among the 535 interacting odorant-receptor pairs, 475 pairs exhibit activation (*´S*_max_ > 0, red), and the remaining pairs show deactivation (*δS*_max_ < 0, cyan). Note that the red and cyan distributions sum to make the gray (all pairs) distribution. The EC_50_ values for the pairs exhibiting activation can be fitted effectively to a Gaussian 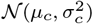 [42] with *μ_c_* = −3.6 and *σ_c_* = 1.1; for all pairs, *μ_c_* = −4.0 and *σ_c_* = 1.8. The skewness of the distribution is likely to be contributed from the extremal (cutoff-based) nature of the measurement, which unavoidably eliminates the weakly interacting (high EC_50_) pairs from the data. The deactivating pairs are characterized with a broader distribution of EC_50_ values. The superstrong binders (EC_50_ < −9) are exclusively contributed by those among the odorant-receptor pairs demonstrating the deactivation. **b**. Odorant-specific differential response for the top five broadly-interacting odorants.

We also determined odorant-specific differential response, *ψ_λ_*(log_10_ *C_O_*), defined as:

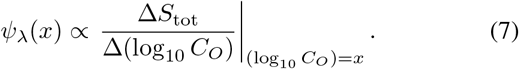

This measures the change of cumulative signal from all *N_r_* receptor types (Δ*S*_tot_) that can be elicited as the log-concentration of the given odorant *λ* increases by Δ(log_10_ *C_O_*). Under an assumption that the odorant *λ* activates a continuous spectrum of receptors, we calculated *ψ_λ_*(log_10_ *C_O_*) for each of the top five broadly-interacting odorants (cis-3-hexen-1-ol, granyl acetate, eugenol, eugenol acetate, butyric acid) that have a sufficiently large number of responsive OR partners (≳ 20). For these broadly-interacting odorants, *ψ_λ_*(log_10_ *C_O_*)’s are sharply peaked around an odorant concentration of 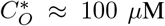 (Fig 3b). Incidentally, 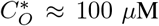 is comparable to the odorant concentration of ~ (10 − 500) × 10^14^ molecules/mL used in the olfactometer measurement [43].

Next, we constructed the distribution of the efficacy, *ρ*(*E*), as a histogram over the ensemble of all odorant-OR pairs. We note that the maximum-entropy distribution of *E*, for an ensemble of odorant-OR pairs characterized with an average efficacy 〈*E*〉, exhibits an exponential functional form *ρ*(*E*) ~ exp(-*E*/〈*E*〉), as was pointed out in [17] to explain the firing rate distribution of the glomeruli. In the case of Mainland *et al*.’s dataset, *ρ*(*E*) was characterized by a sum of two exponential distributions:

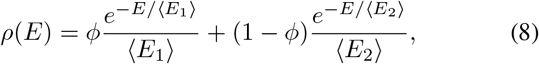

with *φ* = 0.79, and with the two average efficacy values 〈*E*_1_〉 = 1.50 and 〈*E_2_*〉 = 10.06 (Fig 4a). This double-exponential distribution implies that the population of odorant-OR pairs are divided into two subgroups, one characterized with a low efficacy (79% of total population) and the other with a high efficacy (21% population). On the other hand, analysis on the relative amplitude of activation (*δS*_max_) gives rise to qualitatively the same distribution *ρ*(*δS*_max_) as Eq 8 with *φ* = 0.57, 〈*δS*_1,max_〉 = 0.97, and 〈*δS*_2,max_〉 = 5.77. Whether the spiking rates over the population of human ORNs (labeled by their OR subtypes) also constitute two exponential distribution is an interesting question that may be examined by carefully designed experimental measurements.

**Figure 4:**
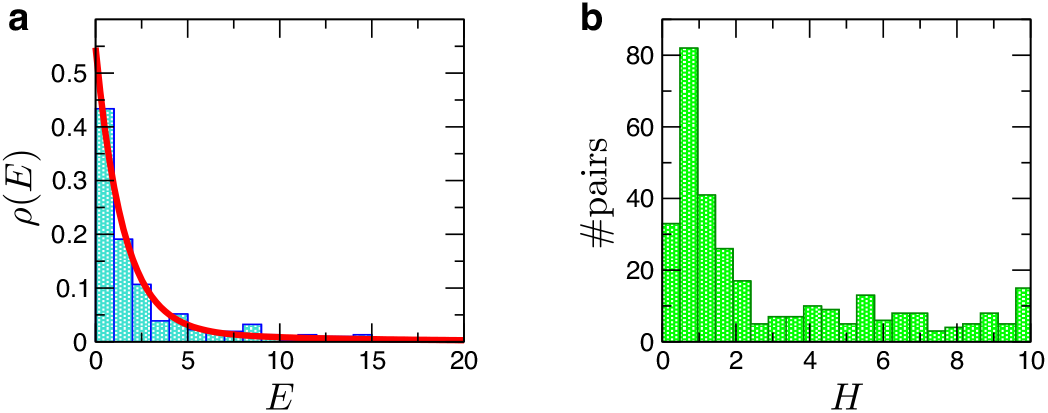
Distributions of efficacy and Hill coefficient. **a**. Probability density of efficacy *ρ*(*E*) and **b**. the histogram of Hill coefficient for the ensemble of all interacting odorant-OR pairs. *ρ*(*E*) was fitted to a sum of two exponentials (Eq 8).

Finally, we report the distribution of Hill coefficients. While a majority of the data are indeed well described with *H* = 1, some are better fitted with larger values of *H* (Fig 4b). Within our two-state activation model, the *phenomenological* value of *H* > 1 could arise from multiple sources (See Supporting Information for details): (i) a cooperative activation of the receptors, which could be linked to the recent studies on the effect of GPCR dimers or higher-order oligomers on the signaling [44, 45, 46, 47, 48, 49]; (ii) the sensitivity amplified in the process of signal cascades accompanying covalent modifications, such as GTP hydrolysis and phosphorylation/dephosphorylation [50, 51]. Given that G-protein pathway is associated with GTP hydrolysis, a scenario of the amplified sensitivity can be considered; (iii) inhomogeneity of the odorant-OR kinetics.

We note that an analytic expression for the activity can still be obtained for either of the foregoing three mechanisms, albeit more convoluted than the Hill equation (Eq 3), and be used to fit the data. Given the expression of the activity, its sensitivity with respect to the variation of control parameter (or the sharpness of transition) can be evaluated by using

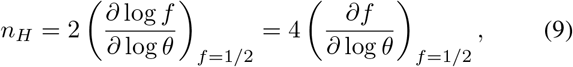

where *f* and *θ* are normalized response and the control parameter, respectively [51]; For the case of Eq 3, *θ* = (*C_O_*/*K*_1/2_) and *f* = *θ^H^*/(1 + *θ^H^*), and it is easy to confirm that *n_H_* = *H*.

### Determination of microscopic rate and binding constants

From Eqs 4-6, we observe that all three kinetic parameters {*E, B, K*_1/2_} depend on the concentration of G-protein (*C_G_*). By eliminating the *C_G_*-dependence from these expressions, we can relate the three parameters in pairs. For example, the efficacy *E* of an odorant-OR pair has a hyperbolic dependence on *B*:

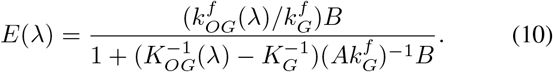

Similarly, *B* is related to *K*_1/2_:

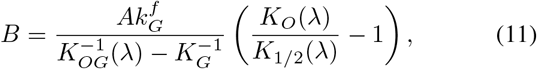

and *E* to *K*_1/2_:

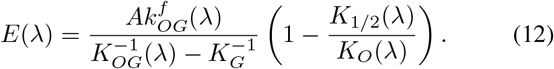

In addition, an interesting triadic relation between the three parameters *E*, *B*, and *K*_1/2_ define a quantity *ω* that can be expressed in terms of the microscopic rates 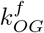, 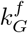, and *K_O_*:

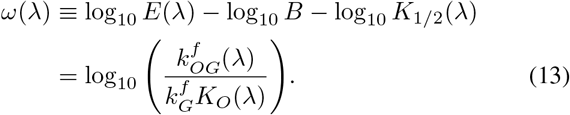

Using Eqs 10-13, we can in principle determine microscopic quantities such as 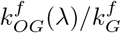, *K_O_*(*λ*), 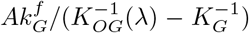, and 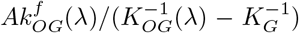 for a given pair of odorant and receptor types.

Here we report the odorant-specific estimates for some of these quantities, averaged over the observed spectrum of receptors that interact with a given odorant *λ*. For example, we obtained 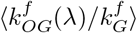 as the amplitude of the *E*(*λ*) versus *B* curve using Eq 10 (Fig 5a). It was then plugged into Eq 13 to estimate 〈log_10_ *K_O_*(*λ*)〉 from the knowledge of 〈*ω*(*λ*)〉, a value that can be obtained from the histogram over all interacting pairs involving this odorant *λ* (Fig 5b). The estimated average values for the top five broadly-interacting odorants are summarized in Table 1.

**Figure 5:**
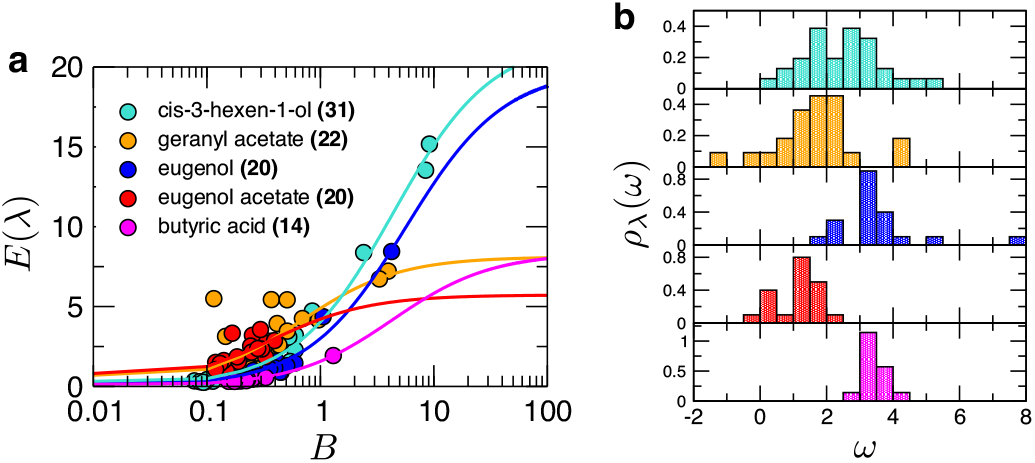
Analysis of the top five broadly-interacting odorants. **a**. Fits of efficacy (*E*) versus basal activity data (*B*) for distinct odorants (*λ*) using Eq 10. The name of each odorant is provided with the number of responding receptors inside the parenthesis. **b**. Distributions of *ω* for the five broadly-interacting odorants.

**Table 1:** OR averaged parameters determined for the top five broadly-interacting odorants. Ratio of rate constants, binding free energy in *k_B_T* unit, and EC_50_ of the top five broadly-interacting odorant, averaged over the multitude of partner receptors.

| Odorant, $\lambda$ | $\langle k_{OG}^f(\lambda)/k_G^f \rangle$ | $\langle \log_{10} K_O(\lambda) \rangle$ | $\langle \log_{10} K_{1/2}(\lambda) \rangle$ |
| --- | --- | --- | --- |
| cis-3-hexen-1-ol | 4.96 | -1.82 | -3.72 |
| geranyl acetate | 12.32 | -0.57 | -4.16 |
| eugenol | 3.53 | -2.91 | -4.41 |
| eugenol acetate | 15.89 | 0.12 | -3.48 |
| butyric acid | 1.93 | -3.13 | -4.14 |

From the kinetic parameters determined here, we make three points that are noteworthy:

i. We find that 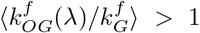 for all the five odorants tested. This is consistent with our expectation that agonist (odorant) binding facilitates the accommodation of G-protein.
ii. *K_O_*(*λ*), the threshold concentration of the odorant *λ* for binding the receptor, is greater than *K*_1/2_(*λ*), the effective threshold for the odorant to eventually elicit the signal (the release of G*α*-subunit). Together with an augmented sensitivity (*H* > 1) discussed above, the relationship of *K_O_* > *K*_1/2_ appears to be a natural outcome of signal cascade in biochemical reactions [50].
iii. Eq 4 with the condition *K_O_* > *K*_1/2_ suggests that *K_OG_* < *K_G_*; the binding of G-proteins to a receptor is stronger when there is an odorant bound to the receptor. This is equivalent to the condition of 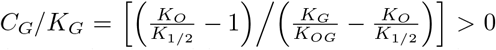, an in-equality that can be derived by rearranging Eq 4. Therefore, it follows that

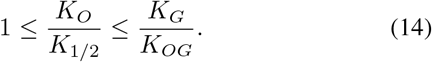

This specifies a necessary condition to be satisfied, within our kinetic model, for the odorant-evoked activity. In other words, for the odorant-dependent activation mechanism to function properly, the *difference* in free energies of G-protein binding for the two cases with and without odorant in the binding site, *ought to* be greater than that between OR activation and odorant binding. The data reported in Table 1 indeed confirm our analysis.

## Discussion

### Non-uniform sensitivity of ORs to odor concentration

Our analysis on Mainland *et al*.’s data [34] provides a new insight into the issue of odor sensitivity. Since qualitatively similar trend is observed in the histogram for the entire ensemble of odorant-OR pairs as shown in Fig 3a, we assume that it effectively represents a general sensitivity profile of any odorant against the pool of human ORs; in other words, we hypothesize that the variable log_10_ *C_O_* displays the identical distribution *ψ*_ens_(log_10_ *C_O_*) as *ψ*_ens_(EC_50_) though *ψ*_ens_ was constructed as a distribution of EC_50_ values. The range of threshold concentration of odorants, at which the olfactory system starts to respond, is finite. To be specific, Fig 3a suggests that *R* = (log *C_O_*)_max_ − (log *C_O_*)_min_ ≈ 2 · (2*σ*_EC_50__) · log_10_ ≈ 16 along the natural logarithmic scale, where (log *C_O_*)_max_ and (log *C_O_*)_min_ denote, respectively, the maximum and minimum values of log *C_O_* that constitute the distribution of EC_50_ and *σ*_EC_50__ is the standard deviation of the distribution *ψ*_ens_(EC_50_). It has been argued previously that the minimal concentration change (Δ*C_O_*) that gives rise to a detectable difference, of at least one bit of the receptor code, is dictated by the Weber ratio Δ*C_O_*/*C_O_* ≈ *R*/*N_r_* ≈ 0.042 [52]. This is to say that, in order for humans to be able to sense a change in odor strength, the concentration difference should exceed 4% in the concentration range between (log *C_O_*)_min_ and (log *C_O_*)_max_. However, this estimate of the Weber ratio was obtained under an assumption that the threshold concentration of odorant is uniformly distributed over the entire range *R* [16, 52].

As shown in Fig 3b, the differential response of receptors against each of the five broadly-interacting odorants displays a non-uniform distribution, *ψ*_λ_(log_10_ *C_O_*), sharply peaked at 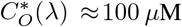. the expected number of ORs that could recognize an odorant at a concentration *C_O_*, denoted *n_O_*, would satisfy

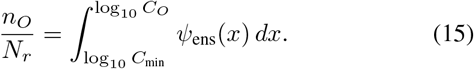

Therefore, the condition for a minimal change in the odorant concentration to activate an additional OR type (Δ*n* ≥ 1) can be expressed using the following inequality:

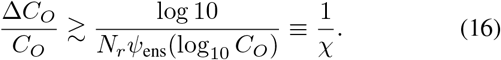

The lower bound for the Weber ratio is minimized at the peak of the sensitivity curve 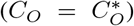, which in turn maximizes the sensitivity *χ* defined above. This implies that the olfactory system is most sensitive at *C_O_* ≈ 100 *μ*M. Using *N_r_* ≈ 330 and *σ*_EC_50__ ≈ 1.5, the minimized value of the Weber ratio is 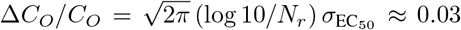; the detectable difference at the sensitivity peak can be as small as 3%.

### Response of the OR cell is effectively binary

According to our analysis of the dose-response data, the sensitivity of OR response is maximized at 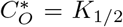. The full width at half maximum (FWHM) of ∂S/∂ (log_10_ *C_O_*) depends on *H* as 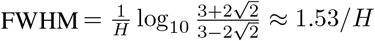. At the level of a single OR, a higher OR sensitivity is attained with a greater *H*.

On the other hand, there is another source of “sensitization” at the level of the ORN, arising from the fact that an ORN is activated when a sufficient number of its ORs are activated together. It is known that a monogenic OR expression (i.e., the expression of only one out of *N_r_*-OR types) is ensured in each ORN, regulated by a feedback mechanism [53, 54, 55, 56]. Therefore, the cellular response of the ORN results from the collective action of *L* ~ 2.5 × 10^4^ receptors of the same type [36]. Even when the activation of individual receptors is stochastic, the collective signal from *L* receptors is much “sharper” at the ensemble level, effectively eliminating the noise in the cellular response. Since the action potential of neuron is switched on and off around a threshold membrane potential 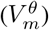, which in turn can be related to a threshold value in the fraction of activated ORs (*ℓ^È^*/*L*), the firing probability of the neuron is effectively binarized as:

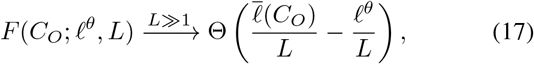

where *Θ*(*z*) = 1 for *z* > 0, and *Θ*(*z*) =0 for *z* ≤ 0 (see Supporting Information for the details of derivation). Because the condition L ≫ 1 eliminates the fluctuations in the copy number of activated receptors, the mapping between the odorant concentration and the activated OR population 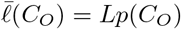 is *deterministic* in every practical sense. The firing probability of the neuron can thus be written as 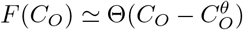, in terms of a threshold concentration 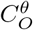; the neuron fires if the odorant concentration is greater than the threshold value 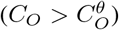.

The effective binarization of electrophysiological signals in the OR cells projects the chemical representation of the odor (response at the sub-cellular level) onto an *N_r_*-bit digital signal (response at the neural level). This *N_r_*-bit information transmitted to the post-synaptic neurons is further processed through the brain circuits for various computational tasks [1, 32, 57]. Although synaptic transmission is still subject to noise, for example due to the stochasticity in the number of vesicles discharged at synapses [58], our argument for the binarization of ORN responses based on the law of large number and activation threshold can still be applied to information transmission that occurs in the upper brain.

### The OR signal is almost fully exploited in odor perception

Putting together our observations so far, the maximal amount of olfactory information that can be encoded into the ensemble of OR cells is *N_r_* bits, corresponding to 2^*N_r_*^ distinct receptor codes. Therefore, the number of distinct odors Ω that can be discriminated by the human olfactory system is upper-bounded as Ω ≲ 2^*N_r_*^. Provided that all natural odors could be represented as a mixture of non-overlapping, equal-intensity odorant components, the ability to discriminate any pair of *m*-component odorant mixtures without ambiguity is subject to the following condition:

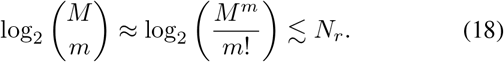

Using the estimated sizes for the human olfactory system, *N_r_* ≈ 330 [26] and *M* ≈ 10^4^ [27, 28], we obtain *m*_max_ ≈ 35 as the maximal size of such odorant mixtures. In other words, the space of all odor mixtures composed of more than *m*_max_ ≈ 35 odorants would exceed the capacity of the human olfactory system. Taking the analogy of the visual system with three receptors (R,G,B), one could further postulate an “olfactory white”, where well-mixed odors with more than *m*_max_ components, each of which elicits an equal intensity response, are indistinguishable from one another [59]. Remarkably, our rough estimate of *m*_max_ « 35 from a simple information-theoretic argument is in reasonable agreement with an experiment [59], where it was shown through an ingenious set of psychophysical tests that well-mixed odors of *m*_max_ ≈ 30 odorant components are indeed indistinguishable to humans.

## Conclusion

Sensing of smell is ultimately a mapping from the molecular information space of odorants to patterns of neuronal activity, which are then perceived as particular kinds of odor in the brain. Understanding the nature of this mapping remains challenging due to factors such as the complexity in the molecular space of odorants, lack of information on the molecular recognition by ORs, and the difficulty of deciphering neuronal signal processing. The present work makes an important step forward in this direction by analyzing the sub-cellular process of olfactory sensing within the ORN cell, at a scale larger than the individual molecular interactions but smaller than the multi-cell signal. Employing a minimal kinetic model for odorant-OR interaction at single molecule scale, we quantified the statistics of interactions between odorants and human ORs, and discussed how the response of individual ORNs can be effectively binarized. The quantitative insights provided in this work can lead to the next level of understanding of human olfaction at multicellular scale.

While the analyses of OR responses in this study are limited to the response of single olfactory receptor at the individual or population level, each odorant is in general recognized by a finite number of multiple ORs. Thus, it is essential to study the combinatorial nature of odor signal processing in a more systematic way based on concrete data and bring the current understanding of olfaction to a systems level. More specifically, since Mainland *et al*. ‘s dose-response data [34] offer such opportunity, we plan to quantify which set of ORs are sampled for a given odorant or odorant mixture and then generate the corresponding receptor code, and also address how discriminable these receptor codes are in the early layer of information processing in the human olfaction. In particular, our model will enable the prediction of receptor responses to mixtures of multiple odorants at possibly different concentrations, which is typical in natural odors. Given that the sensory world through olfactory process is only two synapses away from the cortical neurons [60], addressing these questions will provide better glimpses into the neurobiological principle of signal processing in the human brain.

## Acknowledgments

We thank Leslie Vosshall for helpful comments on the estimated number of monomolecular odorants. SJJ thanks the support from the National Science Foundation (CHE-1362926) and discussion with Kevin Gardner regarding the observation of rigidity of GPCRs when interacting with inverse agonists. We acknowledge the Center for Advanced Computation in KIAS for providing computing resources.

## Supporting Information

### Deviation from the Michaelis-Menten kinetics

The deviation of the odor response from the Michaelis-Menten (MM) form of Eq 2, namely the fitting of Hill curve with *H* ≠ 1, could originate from multiple sources. Here we describe three possible scenarios where we can observe such deviation, and offer relevant quantitative analyses.

#### A cooperative activation of oligomerized receptors

Let us consider the situation where the receptors form a dimer, resulting in two binding sites to which a specific type of odorant can bind. Equilibrium constants for the two binding sites, given as *K*_1_ = [OR][O]/[OR · O_1_] and *K*_2_ = [OR · O_1_][O]/[OR · O_2_], yield the following fractional occupancy of the dimeric complex, which is translated to an odorant concentration (*C_O_* = [*O*])-dependent OR activity:

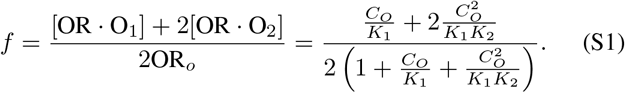

where OR_o_ = [OR] + [OR · O_1_] + [OR · O_2_]. In this case, the Hill coefficient is obtained using Eq 9 as:

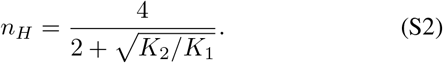

If the two binding sites have positive cooperativity, we have *K*_2_ ≤ *K*_1_, which gives rise to 1 ≤ *n_H_* ≤ 2.

#### The amplification of sensitivity through the signal cascades via GTP hydrolysis

Even in the absence of allosteric cooperativity, a highly sigmoidal, switch-like response can arise from a reversible covalent modification along the signaling pathway. The reversible covalent modification is exemplified by the signaling processes such as phosphorylation/dephosphorylation and GDP/GTP exchange accompanied with GTP hydrolysis, whose effect on the sensitivity of signaling is a well studied issue [61, 62, 51] since the seminal work by Goldbeter and Koshland [50].

In the context of our study, the amount of GDP-bound G-protein (*G_D_*) in response to the stimuli (odorant) defines the olfactory activity. Although our minimal kinetic model for odorant-OR kinetics did not explicitly take into account the effect of GDP/GTP exchange in G-protein and recycling of GDP from GTP hydrolysis, such mechanistic details can modulate the sensitivity of olfactory signaling and consequently make the Hill coefficient deviate from unity. Here we provide an overview of amplified sensitivity through covalent modification by explicitly using the terminologies for the OR signaling.

When OR is in the active form (OR_G_ in Scheme 1), it catalyzes the exchange of G_*D*_ into G_*T*_; on the other hand, the GTPase activating protein (GAP) hydrolyzes the GTP in G_*T*_ to produce G_*D*_ back.

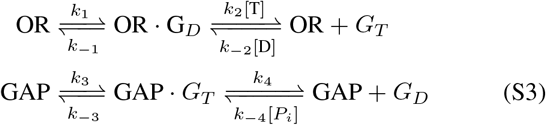

The amount of G_*D*_ in the pool of G-proteins (*G_o_* = [*G_D_*] + [*G_T_*]) is the key quantity that determines the G-protein signaling. The variation of G_*D*_ is given by the difference of the incoming and outgoing currents, *J*_+_ and *J*_−_:

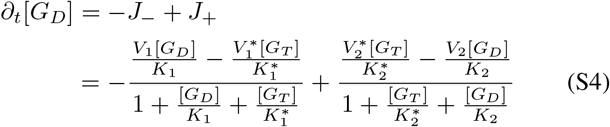

where new notations were introduced for the maximum rates (*V*’s) and Michaelis constants (*K*’s), associated with GDP→GTP exchange in G-protein by 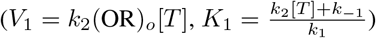; reverse exchange of GTP→ GDP in G-protein 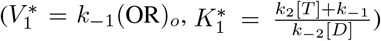; GTP hydrolysis by GAP 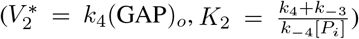; and GTP synthesis 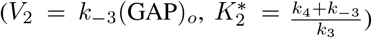. It is useful to define three dimensionless parameters:

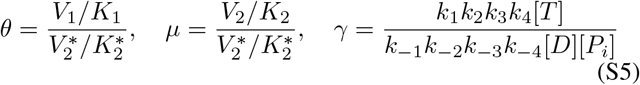

where *γ* is associated with the net non-equilibrium chemical driving force of G-protein signaling pathway in the form of Δ*ψ* = -*k_B_T* log *γ*. Note that Δ*ψ* = 0 (or *γ* =1) at equilibrium.

From the steady state solution *∂_t_*[*G_D_*] = 0 (Eq S4), we can calculate the activity 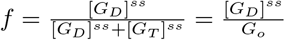 as

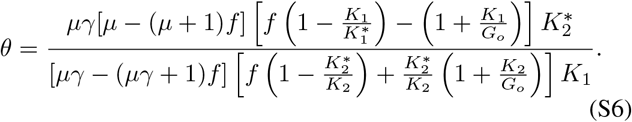

Recall that the neuronal response is directly determined by the amount of *G_D_* (~ *f*), which is in turn dependent on the amount of active OR (~ *θ*). Therefore the sharpness of response *f* (*θ*) is characterized by the effective Hill coefficient, which is once again obtained using Eq 9:

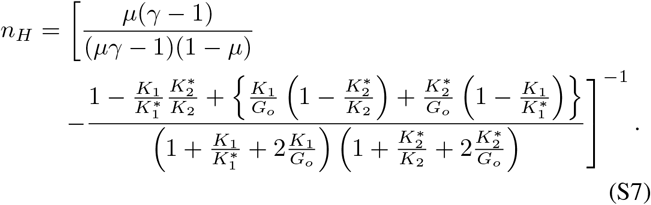

In summary, when GDP/GTP exchange and GTP hydrolysis are explicitly taken into account, the expression of *f* (*θ*) differs from the conventional type of MM expression, and the Hill coefficient evaluated using Eq 9 depends on the parameters *μ*, *γ*, *K*_1,2_, 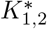, and *G_o_*. Some limiting conditions greatly simplify Eqs S6 and S7:

i. If the affinities of G_*D*_ to OR and G_*T*_ to GAP are sufficiently high 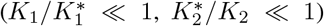 that the overall reaction current associated with the production of G_*D*_ is positive (*J*_+_ − *J*_−_ ≫ 0), and if the G-protein level is below *K*_1_ and 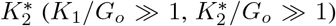, the expressions for *f* and *n_H_* are simplified as:

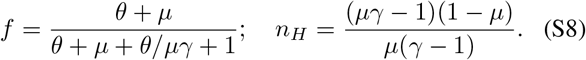
ii. In addition to the aforementioned condition of high affinities of G_*D*_ to OR and G_*T*_ to GAP, if the reversibility of catalytic step is abandoned (*μ* = 0) together with high chemical driving force imposed by a far-from-equilibrium balance of GTP versus GDP (*γ* ≫ 1), the activity *f* is given as

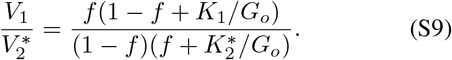

In this case, the Hill coefficient is obtained from 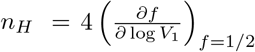 as

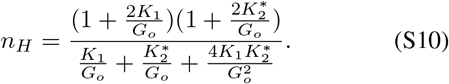

Note that the response is highly sensitized (*n_H_* ≫ 1) for 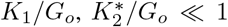. This corresponds to the Goldbeter-Koshland formula for zeroth-order ultrasensitivity [50, 51].

#### Inhomogeneity of the odorant-OR kinetics

Finally, we explore the case when there is inhomogeneity in the parameters for the odorant-OR kinetics. For example, consider the following MM-type hyperbolic activity function:

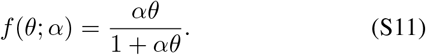

Suppose that the parameter *α* has disorder around its mean value *α*_0_ such that *α* = *α*_0_ + *δα*, |*δα*| ≪ |*α*| where *δα* is a Gaussian random variable satisfying 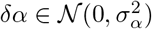, then the above function is approximated up to the second order of *δα* as follows:

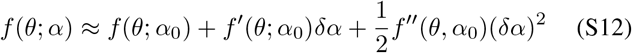

Averaging over the inhomogeneity in *α* with 〈*δα*〉 = 0 and 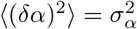 leads to

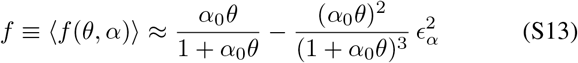

where *ϵ_α_* ≡ *σ_α_*/*α*_0_ ≪ 1. From Eq 9, *n_H_* = 1 for *ϵ_α_* = 0 as expected, and *n_H_* ≈ 0.94 < 1 for *ϵ_α_* = 0.5. The effect is relatively minor compared to the previous two cases, as long as we are in the small fluctuation regime *ϵ_α_* ≪ 1. Nonetheless, the inhomogeneity in parameter is still a possible source of the deviation from MM kinetics. It is interesting to note that the inhomogeneity in kinetic parameter always de-sensitizes the response (*n_H_* ≲ 1).

### Binarized cellular response to odor concentration

Here we outline a simple argument for the effective binarization of cellular responses to odor concentration. Since *p*(*C_O_*) ≡ (*S*(*C_O_*) − *B*)/*δS*_max_ is the activation probability for a single OR at an odorant concentration *C_O_*, the probability of having *ℓ* out of *L* ORs activated is described by a *C_O_*-dependent binomial distribution: 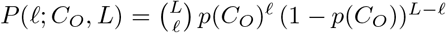. For large number of receptors, satisfying *L*, *Lp*(*C_O_*), *L*(1 -*p*(*C_O_*)) ≫ 1, the binomial distribution is approximated to a normal distribution with a mean 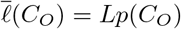 and a variance 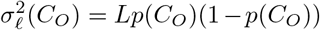. Then, the probability distribution of having *ℓ* out *L* receptors activated is

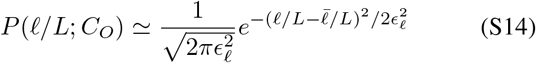

where 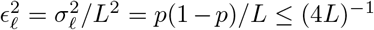. Thus, the fluctuation of the fraction of activated ORs, 〈(*δℓ*/*L*)^2^〉, is suppressed if the population size (*L*) is large. For *L* ~ 2.5 × 10^4^ [36], the size of this fluctuation is limited to less than 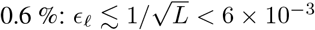.

Meanwhile, the membrane potential V_m_ of a neuron is modulated by changes in the ratio of ion concentrations inside and outside the membranes, which is related to the fraction of open and closed ion channels (or to the fraction of activated and inactivated ORs) (Fig 1a), such that 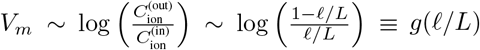 [40, 39]. The first relation is none other than the Nernst equation. Therefore, in the small noise limit one can map CO to 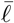 and 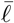 to 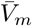, or vice versa, using the monotonic relationships 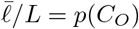 and 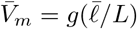, respectively. Note that the generation of a neural spike (action potential) is in general evoked when the membrane potential exceeds a threshold 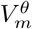 [40, 39]; thus can be effectively related to a threshold in *ℓ*/*L*, or to *C_O_*, such that 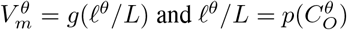 (See Fig S1).

**Figure S1:**
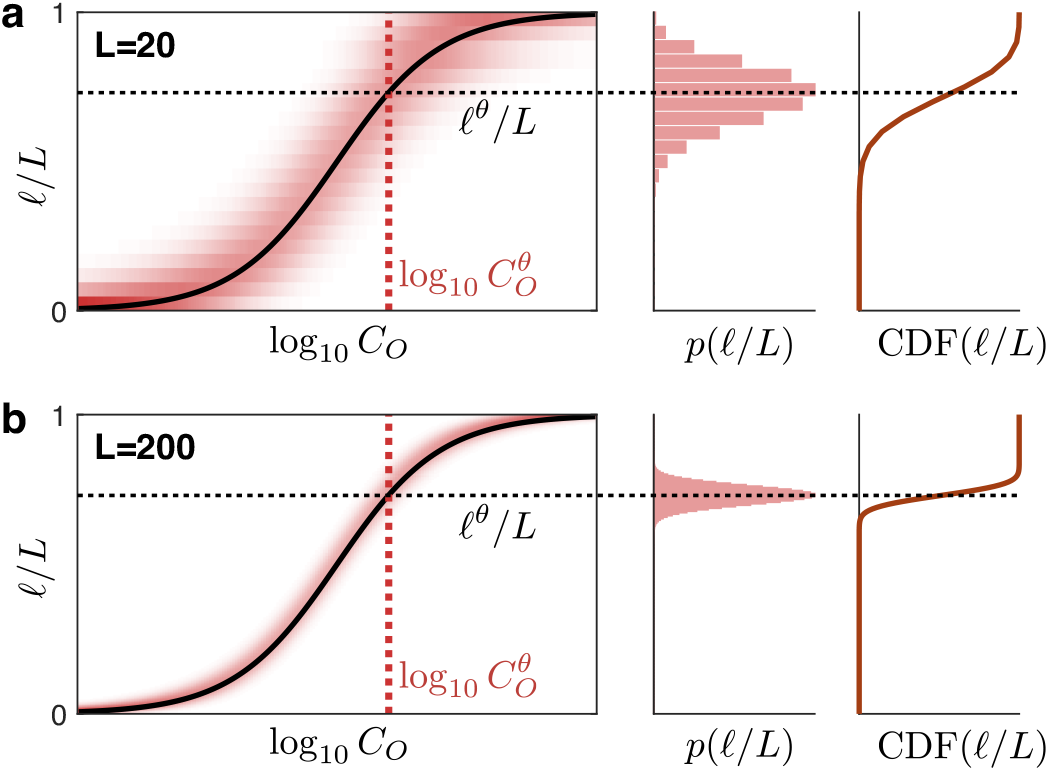
Illustration for the binarized cellular response argument. The sharpness of the response curve depends on the receptor copy number *L* (**a**. *L* = 20, **b**. *L* = 200).

For a given threshold potential, the firing probability of a neuron corresponds to the probability that more than *ℓ^θ^* receptors are activated, and it can be written as:

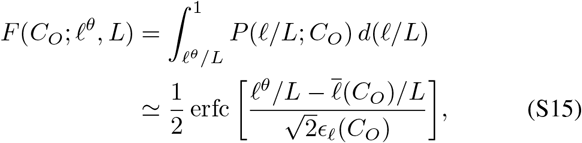

where erfc is the complimentary error function, which is approximated to erfc(*z*) ≃ 1 + sign(*z*) for |*z*| ≫ 2. For large *L*, 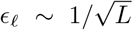, which increases the precision of *ℓ*/*L* for a given *C_O_*. The size of the argument of the erfc in Eq S15 is greater than 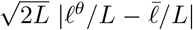; for large L, it is clearly in the |*z*| > 2 regime. Thus, the firing probability can be approximated to the step function as in Eq 17.

**S1 Table. Kinetic parameters determined for all interacting odorant-receptor pairs.** A list of kinetic parameters determined by fitting the dose-response curve from each interacting odorant-receptor pair reported in the dataset of Mainland *et al*. [34] using our model. Table is provided in a comma-separated values (csv) file with 8 columns: olfactory receptor index, odorant index, basal activity (*B*), maximum response (*δS*_max_), efficacy (*E*), EC_50_ (= log_10_(*K*_1_/_2_/[*M*])), Hill coefficient (*H*), and the correlation coefficient of the fit.

**S2 Table. List of odorants tested in the measurement and their chemical types.** A list of all odorants in the dataset of Mainland *et al*. [34] is provided, in a comma-separated values (csv) file with 7 columns. The first five columns are: odorant index, CAS registry number, odorant name, PubChem Compound Identification (CID) number, and SMILES; all as reported in the original dataset [34]. The sixth column shows the chemical type of the odorant. The last column indicates whether this odorant was counted as a strong deactivator in our analysis (Fig S2).

**Figure S2:**
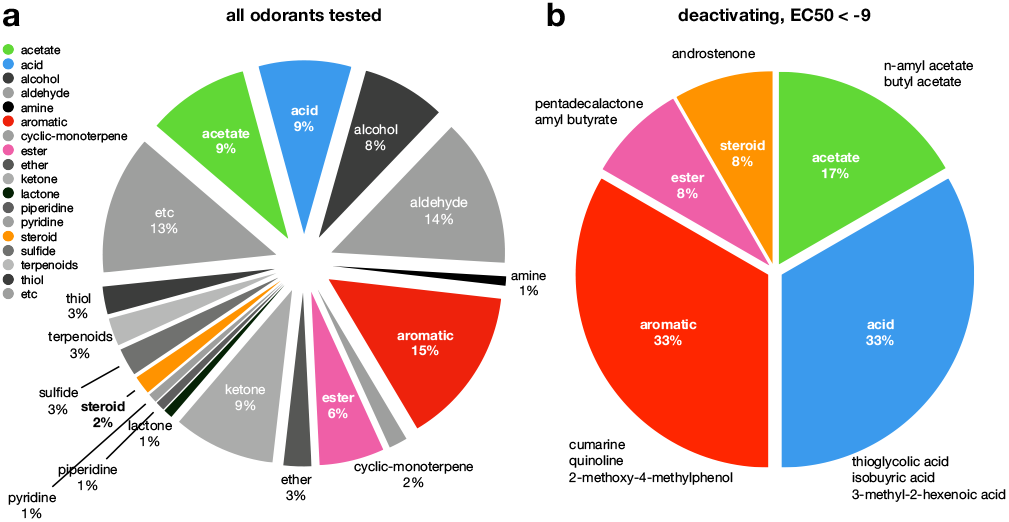
Proportion of odorant types tested in the measurement. **a**. All odorants in dataset from Mainland *et al*. [34] (also see S2 Table). **b**. Odorants with strong affinities (EC_50_ < −9) contributing to the deactivating odorant-OR pairs in Fig 3a.

